# tRNA Biogenesis and Specific Aminoacyl-tRNA Synthetases Regulate Senescence Stability Under the Control of mTOR

**DOI:** 10.1101/2020.04.30.068114

**Authors:** Jordan Guillon, Bertrand Toutain, Coralie Petit, Hugo Coquelet, Cécile Henry, Alice Boissard, Catherine Guette, Olivier Coqueret

## Abstract

Oncogenes or chemotherapy treatments trigger the induction of suppressive pathways such as apoptosis or senescence. Senescence was initially defined as a definitive arrest of cell proliferation but recent results have shown that this mechanism is also associated with cancer progression and chemotherapy resistance. Senescence is therefore much more heterogeneous than initially thought. How this response varies is not really understood, it has been proposed that its outcome relies on the secretome of senescent cells and on the maintenance of their epigenetic marks.

Using experimental models of senescence escape, we now described that the stability of this suppression relies on specific tRNAs and aminoacyl-tRNA synthetases. Following chemotherapy treatment, the DNA binding of the type III RNA polymerase was reduced to prevent tRNA transcription and induce a complete cell cycle arrest. By contrast, during senescence escape, specific tRNAs such as tRNA-Leu-CAA and tRNA-Tyr-GTA were up-regulated. Reducing tRNA transcription appears necessary to control the strength of senescence since RNA pol III inhibition through BRF1 depletion maintained senescence and blocked cell persistence. mTOR inhibition also prevented CIS escape in association with a reduction of tRNA-Leu-CAA and tRNA-Tyr-GTA expression. Further confirming the role of the tRNA-Leu-CAA and tRNA-Tyr-GTA, their corresponding tRNA ligases, YARS and LARS, were necessary for senescence escape. This effect was specific since the CARS ligase had no effect on persistence. YARS and LARS inactivation reduced cell emergence and proteomic analysis indicated that this effect was associated with the down-regulation of E2F1 target genes.

Overall, these findings highlight a new regulation of tRNA biology during senescence and suggest that specific tRNAs and ligases contribute to the strength and heterogeneity of this suppression.

## INTRODUCTION

Senescence induces a definitive proliferative arrest in response to telomere shortening, oncogenes or chemotherapy (1,2). This tumor suppression is most of the time induced by DNA damage and activation of the p53-p21 and p16-Rb pathways. A definitive proliferative arrest is then maintained by the Rb-mediated compaction of proliferative genes within heterochromatin foci or SAHFs (Senescence Associated Heterochromatin Foci) (3). Senescence is also characterized by an increased permeability of the nuclear envelope which allows leakage of chromatin fragments within the cytoplasm. These abnormal fragments are detected by the cGAS-STING DNA sensing pathway which then activates NF-kB and induces the production of a specific secretome known as the SASP (Senescence-Associated Secretory Phenotype). Mainly composed of cytokines and chemokines, this secretome maintains senescence and attracts immune cells which then eliminate the senescent populations (4-7). Thus, through the up-regulation of the p53 and Rb pathways and the activation of immune responses, senescence prevents the propagation of abnormal cells.

Although initially described as a definitive proliferative arrest, we and others have recently shown that some cells can escape senescence, indicating that distinct stages of light and deep senescence should be distinguished (2,7-9). This heterogeneity can be explained by a variable expression of p16 which is necessary for senescence maintenance. The stability of senescence also implies that the compaction of proliferative genes is maintained within the SAHFs. Recent results have reported that H3K9Me3 repressive marks can be removed by the JMJD2C and LSD1 demethylases and that this induces senescence escape (10). The HIRA histone chaperone also plays a key role in the deposition of the histones H3.3 and H4 into the chromatin. Its down-regulation allows senescence escape, indicating again that the stability of epigenetic marks plays a key role in the maintenance of the suppressive arrest. The dynamic nature of senescence is also explained by the variability of the SASP. Initially described as beneficial, several studies have reported that its composition varies and that this secretome can also enhance inflammation or tumor progression (11-13). The reason for this variability is not really understood. Cancer cell lines also represent an interesting model of an incomplete senescence response. We and others have shown that this suppression functions as an adaptive mechanism in response to chemotherapy-induced senescence (CIS) (7,9). We have described that cancer cells can escape senescence and emerge as more transformed cells that resist anoikis and are more invasive (14-17). In addition, we have also shown that cells having an incomplete senescence response are characterized by a reduced expression of CD47 (17). As previously proposed (18,19), this indicates that senescent populations are heterogeneous and can be identified by cell surface receptors.

Taken together, these studies indicate that senescence is much more dynamic than initially thought and that a better characterization of these arrested cells is necessary. At the single cell level, any modification of epigenetic marks, of the SASP composition or any variability of the oncogenic background will generate heterogeneous populations. In this study, we pursued our experiments with the aim of characterizing the signaling pathways involved in the maintenance of senescence. During the initial steps of this suppressive arrest, we describe that the transcription of the type III RNA polymerase is down-regulated and that tRNA synthesis is then reactivated during senescence escape. Depending on the experimental model, results showed that the expression of specific tRNAs was reactivated, such as the tRNA^Leu-^CAA and tRNA^Tyr-^GTA in emergent colorectal cells or breast organoids. In addition, specific aminoacyl-tRNA synthetases (ARS) such as the Leucyl-and Tyrosyl-tRNA ligases were necessary for senescence escape. Our results also indicate that this deregulation of tRNA synthesis led to the activation of the Unfolded Protein Response (UPR) and that this tRNA-mediated ER stress is resolved by mTOR to allow senescence escape.

Altogether, these results indicate that specific pools of tRNAs or ARSs regulate the outcome of this suppressive response and of chemotherapy. We propose that different types of senescence, replicative, oncogenic or mediated by chemotherapy might lead to the expression of different pools of tRNA and ARSs. This could partly explain the variability of the SASP and the deleterious effects of senescence in response to treatment.

## RESULTS

### mTOR activity is necessary for CIS escape

In breast and colorectal cell lines that do not express p16INK4, we have previously reported that p21 maintains senescence and that its down-regulation allows CIS escape (17). To confirm this observation, senescence was induced with sn38 for 96hr, cells were then transfected with control siRNA or siRNA directed against p21 and emergence was evaluated after 10 days (Figure 1A). Western blot analysis confirmed the down-regulation of the cell cycle inhibitor and showed a concomitant up-regulation of cyclin A and of phospho-Rb (Figure 1B). As we have shown (17), this led to a significant increase in CIS escape (Figure 1C). This effect was observed in LS174T colorectal cells and in MCF7 breast cells (Supplementary Figure 1A). We then used mass spectrometry to analyze the signaling pathways involved in CIS escape. To this end, p21 was inactivated in senescent cells, extracts were collected after 48 hr and analyzed by SWATH-MS approaches as we recently described (20). GSEA analysis indicated that Myc and E2Fs proliferative pathways were reactivated as expected. Interestingly, a significant up-regulation of mTOR signaling was also detected (Figure 1D). We focused on this kinase since it plays a key role during senescence (21-25). Western blot analysis confirmed that mTOR was activated in LS174T and MCF7 cells following p21 inhibition, as shown by the phosphorylation of the S6 ribosomal protein, one of its main targets (Figure 1E). We then asked if mTOR was involved in CIS escape by treating the cells at the beginning of emergence with torin-1 and rapamycin, two common drugs used to inactivate this kinase. These inhibitors significantly blocked CIS escape despite p21 inactivation (Figure 1F). Note that low doses of Rapamycin (5 nM) and Torin-1 (15nM) were used to reduce the inhibitory effect on cell proliferation. Rb phosphorylation or cyclin A expression were not inhibited by the two drugs, excluding an unrelated effect on cell cycle progression (Figure 1G). In these conditions, emergent cells were more sensitive to mTOR inhibition than untreated, growing cells (Supplementary Figure 1B). We then determined if mTOR was also involved in spontaneous CIS escape, in the absence of p21 manipulation. Western blot analysis indicated that the kinase activity was increased at the early steps of emergence (Figure 1H). In this condition, Torin-1 or rapamycin inhibited its activity and significantly reduced CIS escape, both in LS174T and MCF7 cells (Figure 1H and I).

**Figure 1:**
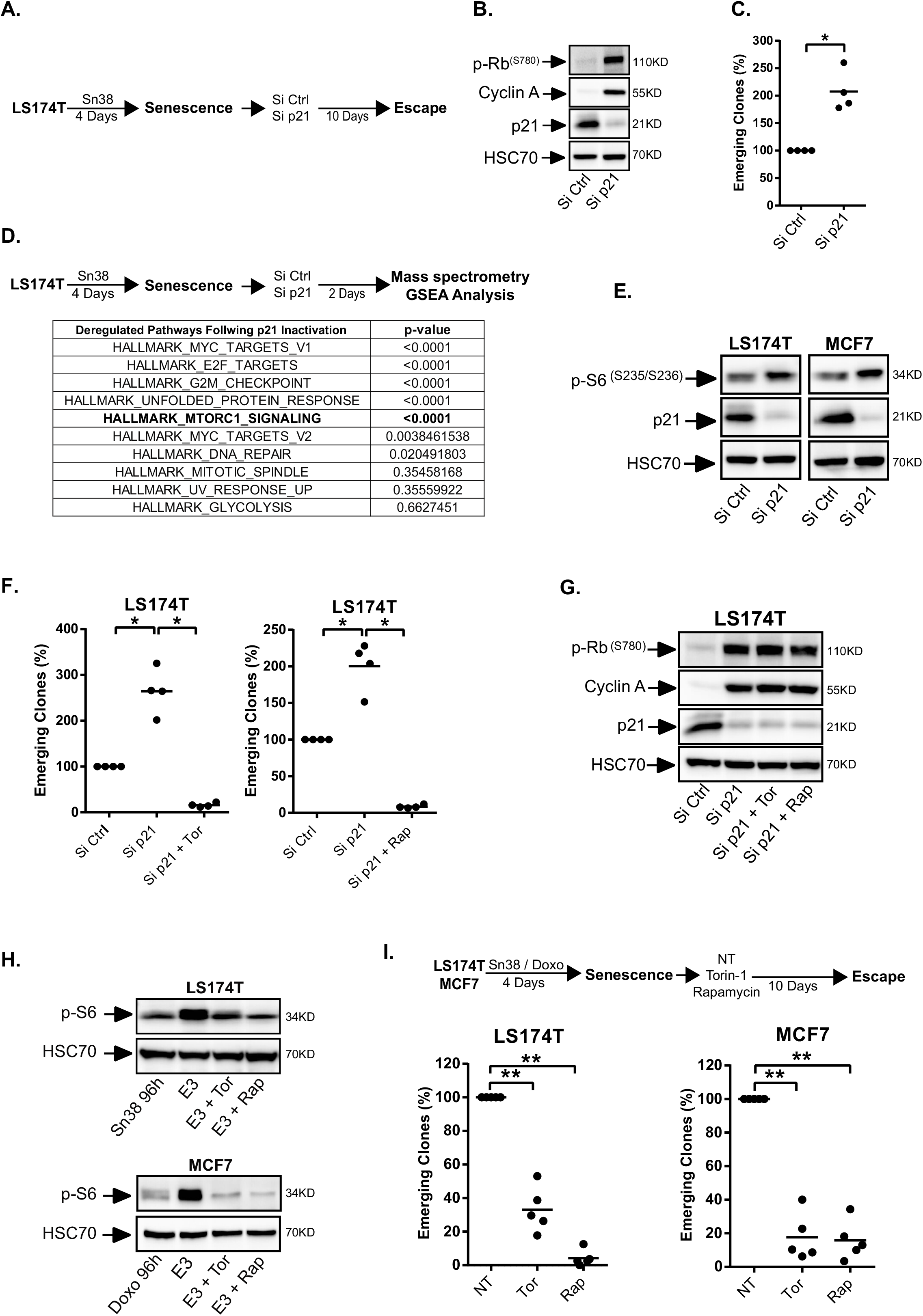
mTOR is necessary for senescence escape. A. Senescence was induced by treating LS174T cells with sn38 as indicated. Cells were then washed with PBS and transfected with a control siRNA or a siRNA directed against p21 for 24 hr and senescence escape was generated by adding 10% FBS. B. LS174T cells were treated as above and cell extracts were recovered 2 days after p21 depletion. The expression of the indicated proteins was analyzed by western blot (n=3). C. Number of emerging clones analyzed after p21 inactivation (n=4, Kolmogorov-Smirnov test * = p<0.05). D. Senescent cells were transfected with a control siRNA or a siRNA directed against p21. Cell extracts were analyzed by SWATH quantitative proteomics and GSEA analysis 2 days after p21 depletion (n=3). Note that this table and experiments are the same as Figure 2A. E. Validation of mTORCI activation by western blot 2 days after p21 inactivation in LS174T and MCF7 cells (n=3). F and G. Senescent LS174T cells were transfected with a control siRNA or a siRNA directed against p21 for 24 hr. Cells were then stimulated with 10 % FBS in the presence or absence of mTOR inhibitors (Rapamycin: 5nM, Torin-1: 15nM). The number of emerging clones was evaluated after 10 days (F, n=4, Kolmogorov-Smirnov test *= p<0.05). Cell extracts were recovered after 2 days and the expression of the indicated proteins was analyzed by western blot (G, n=3). H. Senescent cells were generated as above and emergence was induced by adding 10% FBS. mTORCI activation was analyzed by western blot after 3 days (noted E3 on the figure, n=3 for MCF7, 2 for LS174T). I. Following senescence induction, cells were stimulated with 10% FBS in the presence or absence of mTOR inhibitors. The number of emerging clones was evaluated 10 days later (n=5, Kolmogorov-Smirnov test ** = p<0.01).

Altogether these observations indicate mTOR is necessary for CIS escape, either during spontaneous emergence or following p21 inactivation.

### mTOR promotes CIS escape under proteic stress conditions

The mass spectrometry analysis also detected a significant deregulation of the Unfolded Protein Response (UPR) when cell emergence was induced by p21 inactivation (Figure 2A). This was confirmed during spontaneous CIS escape, as evidenced by an increased expression of Bip and Chop, two main sensors of ER stress signaling. Note that this was detected in emergent MCF7 cells but not in LS174T cells, suggesting that the two cell lines resolved this stress differently (Figure 2B). This suggested that cell emergence might be associated with the ability to resolve this proteic stress. To test this hypothesis, we used tunicamycin or thapsigargin, two well known inducers of the UPR pathway. When added on senescent cells, these two drugs significantly blocked CIS escape. Bip and ChOP expressions were increased as expected (Figure 2C and supplementary Figure 1C). Since mTOR inhibition prevented cell emergence, we then asked if this was related to an abnormal activation of this stress pathway. A significant increase of Bip expression was detected during emergence when MCF7 senescent cells were treated with Torin-1 or rapamycin (Figure 2D). This effect was less evident in LS174T cells, again suggesting that the colorectal cell line might resolve the proteic stress more efficiently. To extend this observation, we activated mTOR using a lentivirus encoding an shRNA directed against TSC2. As an inhibitor of the Reb GTPase, TSC2 is one of the main repressors of the kinase (26). Cells were infected after CIS induction, and senescent cells were treated with tunicamycin to increase the UPR stress and block emergence as described Figure 2C. Results showed that mTOR activation allowed CIS escape despite the increased proteotoxic stress generated by the drug. This effect was observed in MCF7 cells but also in LS174T cells (Figure 2E-F).

**Figure 2:**
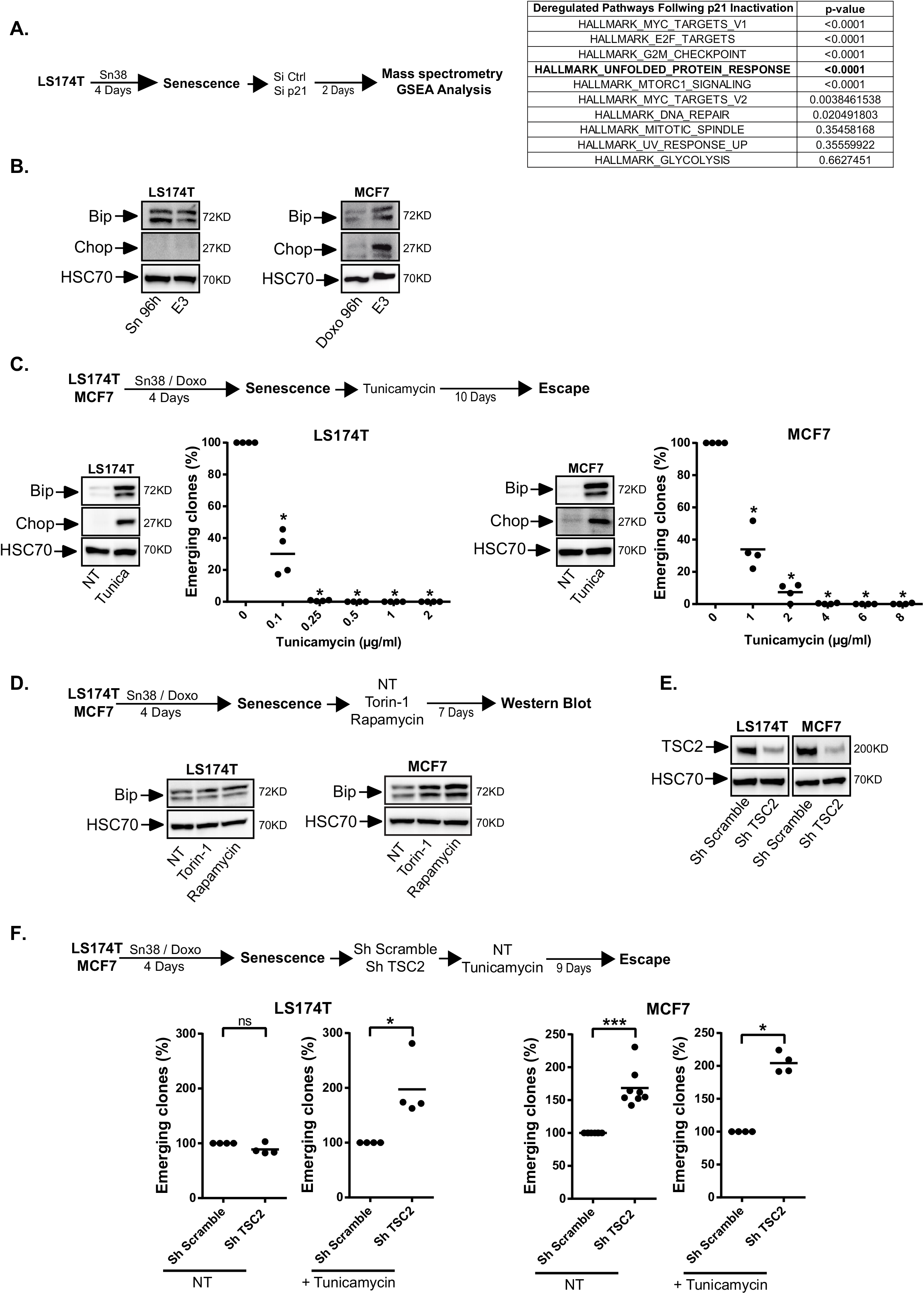
mTOR allows senescence escape in the presence of an ER stress. A. Senescent cells were transfected with a control siRNA or a siRNA directed against p21. Cell extracts were analyzed by SWATH quantitative proteomics and GSEA analysis 2 days after p21 depletion (n=3). Note that this table and experiments are the same as Figure 1D. B. Analysis of the expression of the UPR markers in LS174T and MCF7 senescent cells after 3 days of emergence (noted E3 on the figure, n=3). C. Senescent MCF7 and LS174T cells were treated with increasing concentrations of tunicamycin as indicated and the number of emerging clones was evaluated 10 days later (n=4, Kolmogorov-Smirnov test, * = p<0.05). UPR induction was validated by Western blot 24 hr after the treatment of senescent cells with the lowest concentration of tunicamycin (LS174T: 0.1 µg/ ml, n=2; MCF7 :1 µg/ml, n=2). D. After senescence induction, cells were stimulated with 10% FBS in the presence or absence of mTOR inhibitors. Cell extracts were recovered 7 days later and the expression of Bip was analyzed by western blot (n=3). E, F. Following senescence induction, MCF7 and LS174T cells were transduced with a control shRNA or a shRNA directed against TSC2. TSC2 inhibition was validated by western blot 2 days after transduction (E. LS174T n=2, MCF7 n=1). F: Cells were then washed and treated or not with tunicamycin and emergence was then evaluated as above (n=4 to 8, Kolmogorov-Smirnov test, * = p<0.05 *** = p<0.001).

Together, these results indicate that a high UPR stress prevents CIS escape and that mTOR promotes cell emergence when this stress is increased.

### mTOR Regulates tRNA biogenesis during CIS escape

We then tried to understand the link between CIS escape, mTOR and the ER stress. In LS174T cells, we noticed that rapamycin and torin-1 always reduced the total RNA concentration during senescence escape. Conversely, when TSC2 was inactivated in MCF7 cells, this led to an increased RNA concentration (Figure 3A). Since mTOR is a global activator of tRNA transcription in cancer cells (27), we also determined if this was also the case during CIS escape. Results presented Figure 3B indicate that the expression of several tRNAs was reduced when senescent LS174T cells were treated with Torin-1 or rapamycin. The tRNA^Tyr-^GTA and the tRNA^Leu-^CAA were more significantly down-regulated than the other tRNAs tested. Conversely, when mTOR was activated by the depletion of TSC2 in emergent MCF7 cells, we observed a more significant up-regulation of the tRNA^Tyr-^GTA and tRNA^Leu-^CAA as compared to the other tested tRNAs (Figure 3C). mTOR inhibition also reduced the expression of the 5S rRNA but did not affect RNA polymerase I targets such as the pre45S and 18S rRNAs.

**Figure 3:**
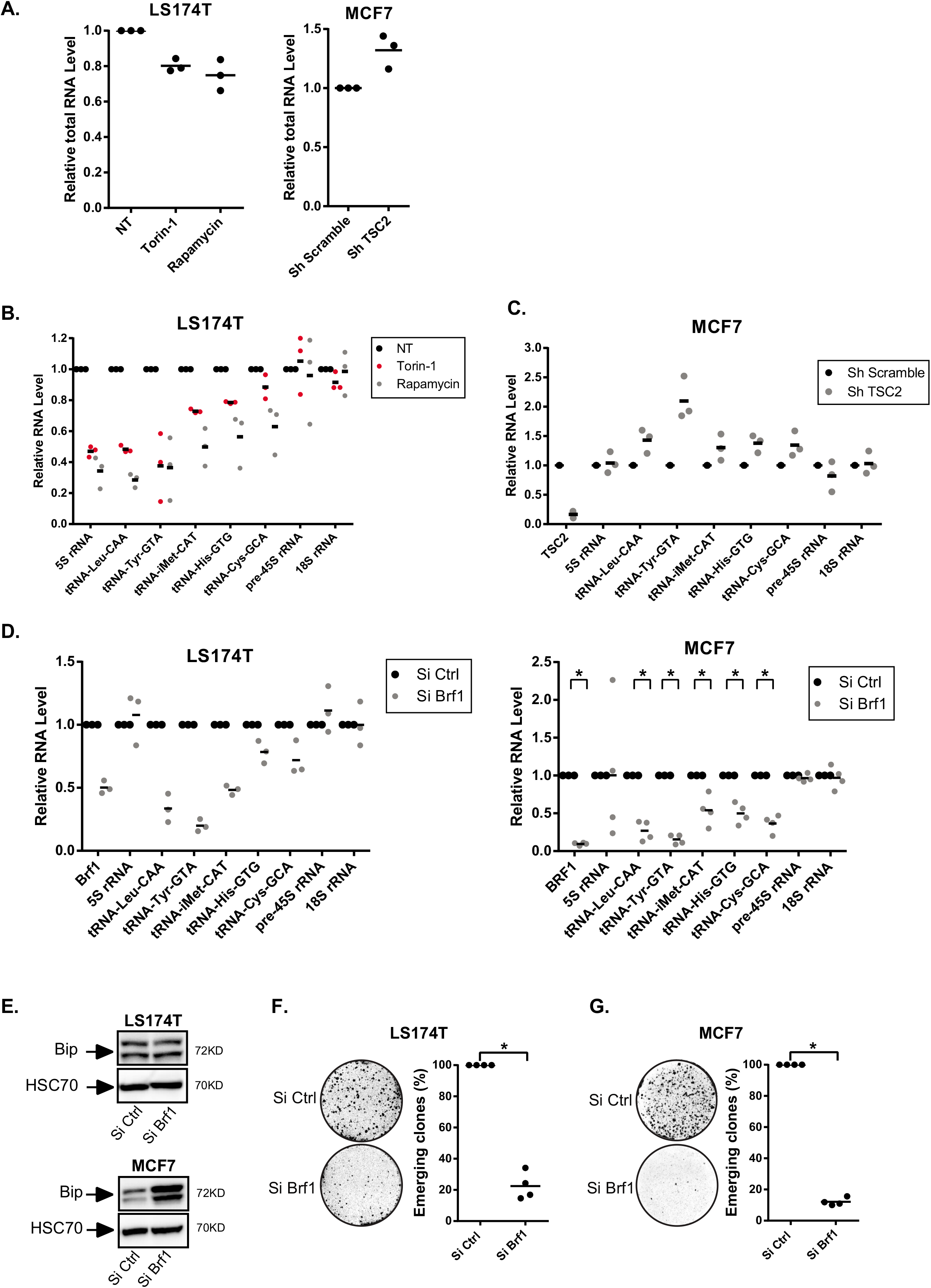
mTOR regulates RNA Pol III activity during CIS escape. A. Quantification of total RNA level in senescent LS174T cells two days after mTOR inhibition (left, n=3) or in senescent MCF7 cells two days after TSC2 inhibition (right, n=3). B. Senescent LS174T cells were treated with mTOR inhibitors, and the relative expression of the indicated RNA was analyzed by RT-QPCR after 7 days (n=3). C. Analysis of the expression of the indicated RNAs 2 days after TSC2 depletion in MCF7 senescent cells by RT-QPCR (n=3). D. Senescent LS174T and MCF7 cells were transfected with a control siRNA or a siRNA directed against Brf1 during 24 hr and then washed with PBS and stimulated with fresh media. Cell extracts were recovered 2 days after Brf1 depletion and the expression of the indicated RNAs was analyzed by RT-QPCR (LS174T: n=3; MCF7: n=4, Kolmogorov-Smirnov test, * = p<0.05). E. Analysis of Bip expression by western blot 7 days after BRF1 depletion in LS174T and MCF7 senescent cells (n=3). F, G. Senescent LS174T and MCF7 cells were transfected with a control siRNA or a siRNA directed against against Brf1 during 24 hr and then washed with PBS and stimulated with fresh media. Nine days after Brf1 inactivation, the number of emerging clones was analyzed (n=4, Kolmogorov-Smirnov test, * = p<0.05).

In light of these results, we then determined if a global inhibition of tRNA synthesis could lead to an increased ER stress. To this end, we inactivated BRF1, the main RNA Pol III activator, in senescent cells. As expected, this reduced tRNA expression in MCF7 and LS174T cells (Figure 3D). This inactivation led to a significant induction of the Bip sensor in MCF7 cells (Figure 3E). This was not observed in LS174T cells, again indicating that these cells may tolerate a higher level of protein stress. When BRF1 was down-regulated, a significant inhibition of senescence escape was observed in the two cell lines (Figure 3F and G). After 7 days of emergence and cell fixation, a microscopic examination allowed the visualization of emergent, dividing, white clones in the middle of blue senescent cells identified by SA-ß-galactosidase staining. These dividing, white clones were completely absent when BRF1 was inactivated, indicating that reducing the activity of RNA pol III allowed the maintenance of senescence (Supplementary Figure 1D).

Altogether, these results indicate that mTOR inhibition reduced tRNA expression and in particular the tRNA^Tyr-^GTA and the tRNA^Leu-^CAA during CIS escape. This led to an increased ER stress indicating that mTOR and RNA Pol III activity are necessary to tolerate the proteic stress during cell emergence.

### Regulation of tRNA expression during senescence and cell emergence

These observations suggested that an up-regulation of tRNA pathways could be necessary to promote CIS escape. To test this hypothesis, we first analyzed tRNA expression in senescent and emergent cells. RT-QPCR results indicated that tRNA expression was down-regulated in senescent cells, both in LS174T and MCF7 cells (Figure 4A and B). In contrast, the expression of the 18S and pre-45S ribosomal RNAs was not significantly modified, further indicating that the activity of RNA pol I was not affected under these experimental conditions. Using chromatin immunoprecipitation experiments, we also observed that the type III RNA polymerase was recruited to the tRNA promoters in growing cells but that its binding was significantly reduced in senescent populations (Figure 4C). These observations suggested to us that this reduction of RNA pol III activity might be a necessary step of senescence induction. To test this hypothesis, we inactivated MAF1 in growing cells and then induced senescence in LS174T or MCF7 cells. MAF1 is the main repressor of RNA Pol III and its down-regulation enhances tRNA transcription (27-29). As presented Figure 4D, the down-regulation of MAF1 led to a significant increase of CIS escape. This effect was specific since MAF1 inactivation did not significantly modify the proliferation of non-treated, growing cells (Supplementary Figure 2A). Thus, a reduction of RNA Pol III transcription during the early step of senescence is necessary to allow a complete proliferative arrest.

We then determined if tRNA expression was up-regulated during CIS escape. A significant increase of the tRNA^Leu-^CAA and tRNA^Tyr-^GTA tRNAs was observed in emergent LS174T cells. The same effect was observed in MCF7 cells where several tRNAs were reactivated (Figure 4E and F). Note that all tRNAs were not up-regulated in emergent LS174T cells and that the tRNA^Cys^ was not significantly modified in both cell lines. The expression of the 45S and 18S rRNAs was also not modified. To determine if this effect was related to senescence or to a consequence of chemotherapy treatment and cell cycle arrest, we repeated these experiments in MCF7 cells treated with a lower dose of doxorubicin. In this condition, cell cycle arrest was induced as observed during the initial step of CIS induction (Supplementary Figure 2B). In this case, no significant inhibition of tRNA transcription was observed and a slight and general reactivation was observed 3 days after the removal of doxorubicin (Supplementary Figure 2C and D).

**Figure 4:**
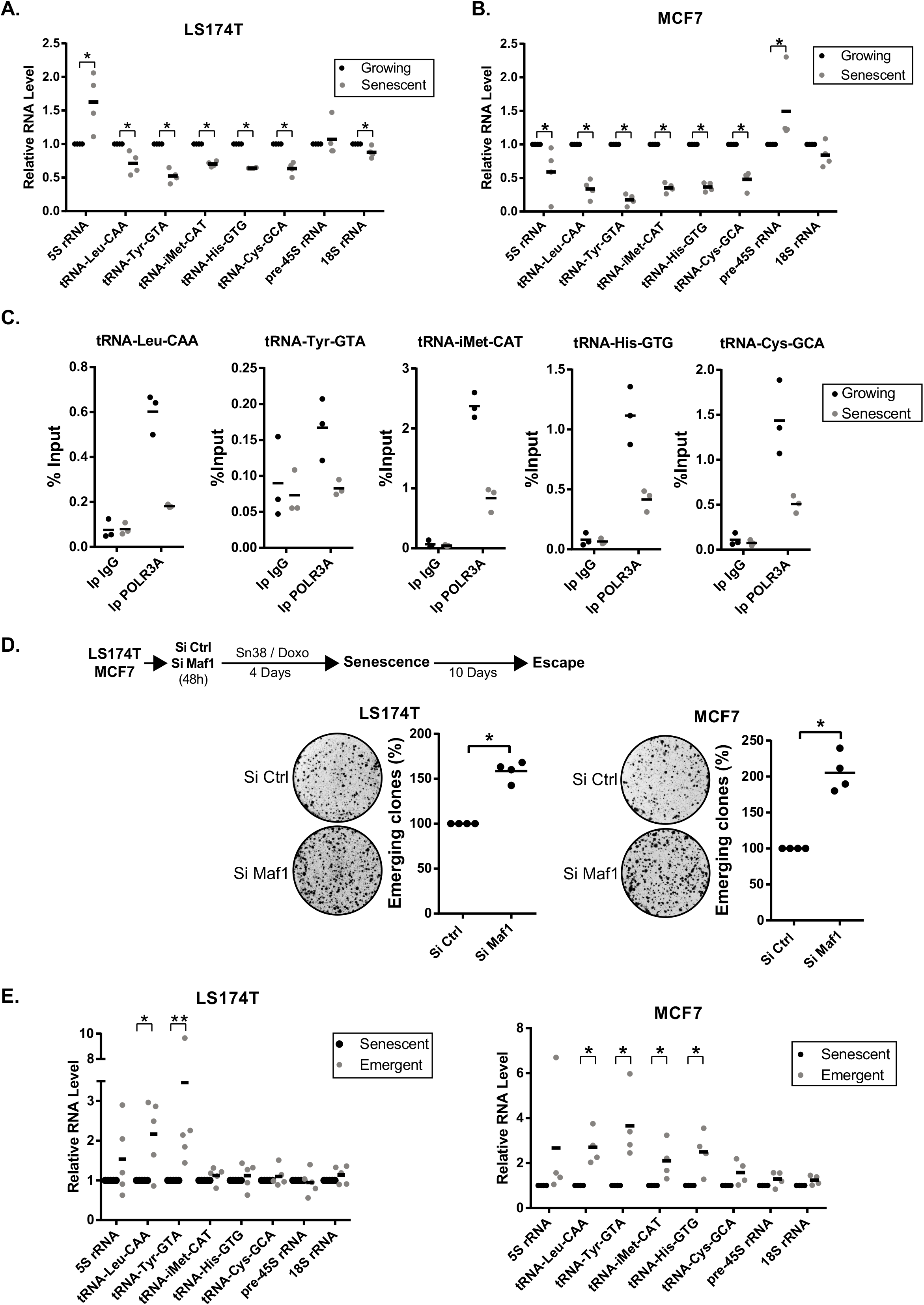
Regulation of tRNA synthesis during senescence and CIS escape. A, B. Expression of the indicated RNAs in growing or senescent cells (LS174T n=4, MCF7 n=4, Kolmogorov-Smirnov test, * = p<0.05). C. ChIP analysis of the binding of the type III RNA polymerase (POLR3A) in senescent or growing LS174T cells (n=3). D. LS174T and MCF7 were transfected with a control siRNA or a siRNA directed against Maf1 48h prior to chemotherapeutic treatment. After senescence induction, cells were washed with PBS and stimulated with fresh media. Ten days later, the number of emerging clones was analyzed (n=4, Kolmogorov-Smirnov test, * = p<0.05). E. Analysis of the expression of the indicated RNAs in senescent or emergent cells (after 7 days, LS174T n=5, MCF7, n=4, Kolmogorov-Smirnov test, * = p<0.05, **= p<0.01).

To extend these observations, we then determined if tRNA transcription was also reactivated in patients-derived organoids (PDO, see (30)) that escape chemotherapy (see a representative image Figure 5A). PDOs isolated from two different patients were treated with doxorubicin and pilot experiments indicated that a proliferative arrest was induced after two sequential drug treatments for 96 hr. As these cells grew in three dimensions, the SA-ß-galactosidase staining was constitutively positive within the inner part of the spheroid, precluding the use of this marker. RT-QPCR experiments showed that proliferative genes such as mcm2 or cdc25A were down-regulated, together with an up-regulation of IL-8 as an illustrative member of the SASP (Figure 5B). P21 expression was also increased in the two PDOs but this was not the case of p16, probably because this gene was already inactivated. After chemotherapy treatment, some PDOs restarted proliferation and this was illustrated after 10 days by the reactivation of mcm2 and cdc25A and the down-regulation of p21 (Figure 5C). Results presented in Figure 5D indicate that the tRNA^Leu-^CAA and tRNA^Tyr-^GTA were also up-regulated. This was not the case for the other tRNAs and ribosomal RNAs tested.

**Figure 5:**
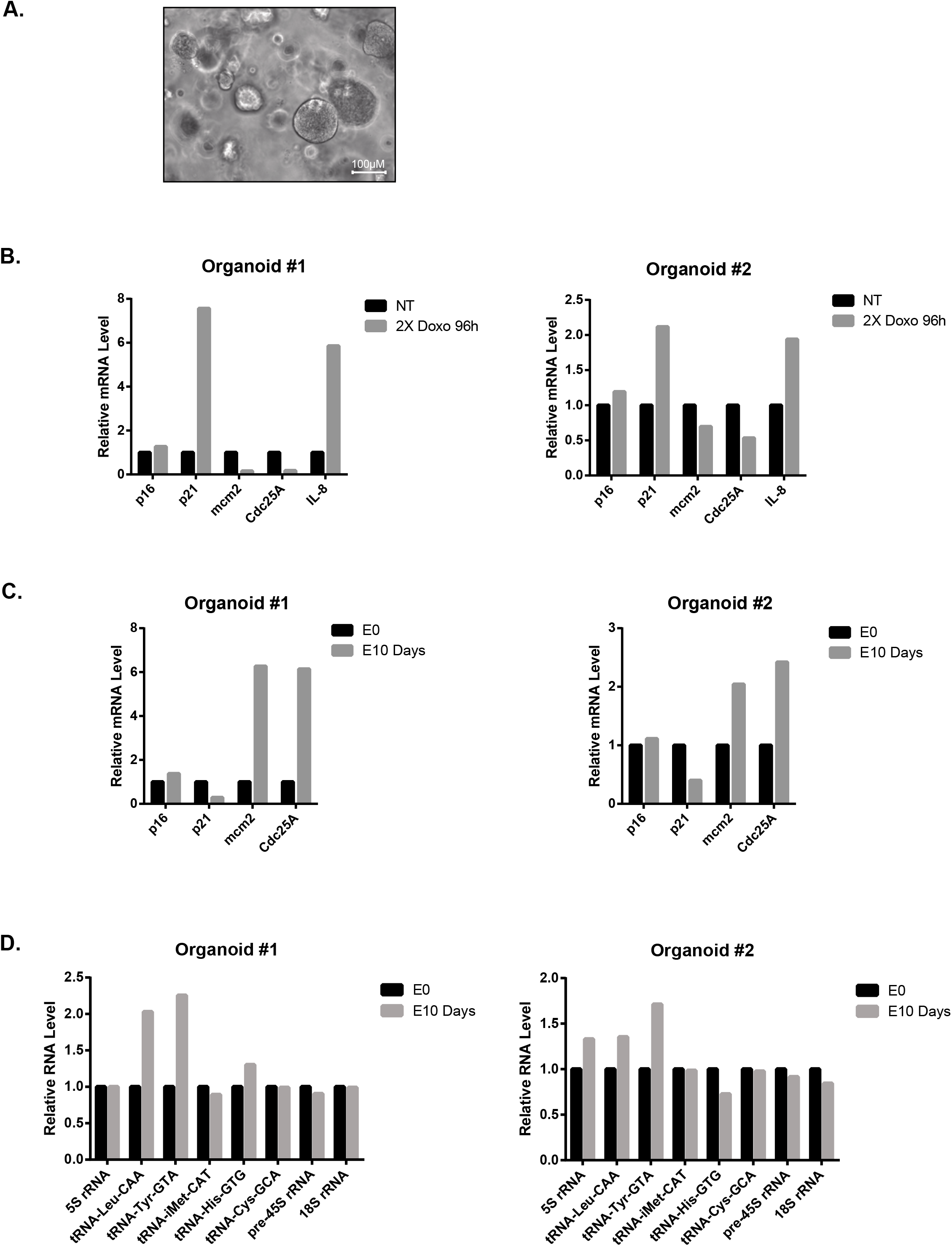
Up-regulation of the tRNA^Leu^-CAA and tRNA^Tyr^-GTA in breast organoids that escape chemotherapy. A. Representative image of breast cancer organoids. B. Analysis by RT-QPCR of proliferative and senescence markers of two breast tumor organoids treated or not with 2 cycles of 96 hr of Doxorubicin (All experiments were performed on two organoids coming from two patients, #1: Doxorubicin 25 ng/ml, #2: Doxorubicin 50ng/ml). C, D. Analysis of the expression of the indicated RNAs on PDOs at the end of the doxorubicin treatment (E0) and 10 days later (E10).

Therefore, in two different conditions of treatment escape, we observed an up-regulation of different tRNAs such as the tRNA^Leu-^CAA and tRNA^Tyr-^GTA when cancer cells resumed proliferation following chemotherapy treatment.

### The LARS and YARS tRNA ligases promote CIS escape

These results suggested to us that CIS escape might be induced either by a global reactivation of RNA pol III activity or by the over-expression of specific tRNAs. One could first speculate that a general activation of tRNA pathways promotes CIS escape by simply increasing protein synthesis. To test this hypothesis, we inactivated MAF1, the RNA Pol III repressor, when senescence was established. This led to a general increase of tRNA expression, both in LS174T and MCF7 cells (Figure 6A). Surprisingly, a weak increase of CIS escape was detected in LS174T cells but no significant effect was observed in the breast cell line (Figure 6B). Thus, a general increase of tRNA expression in senescent cells was not sufficient to induce CIS escape.

**Figure 6:**
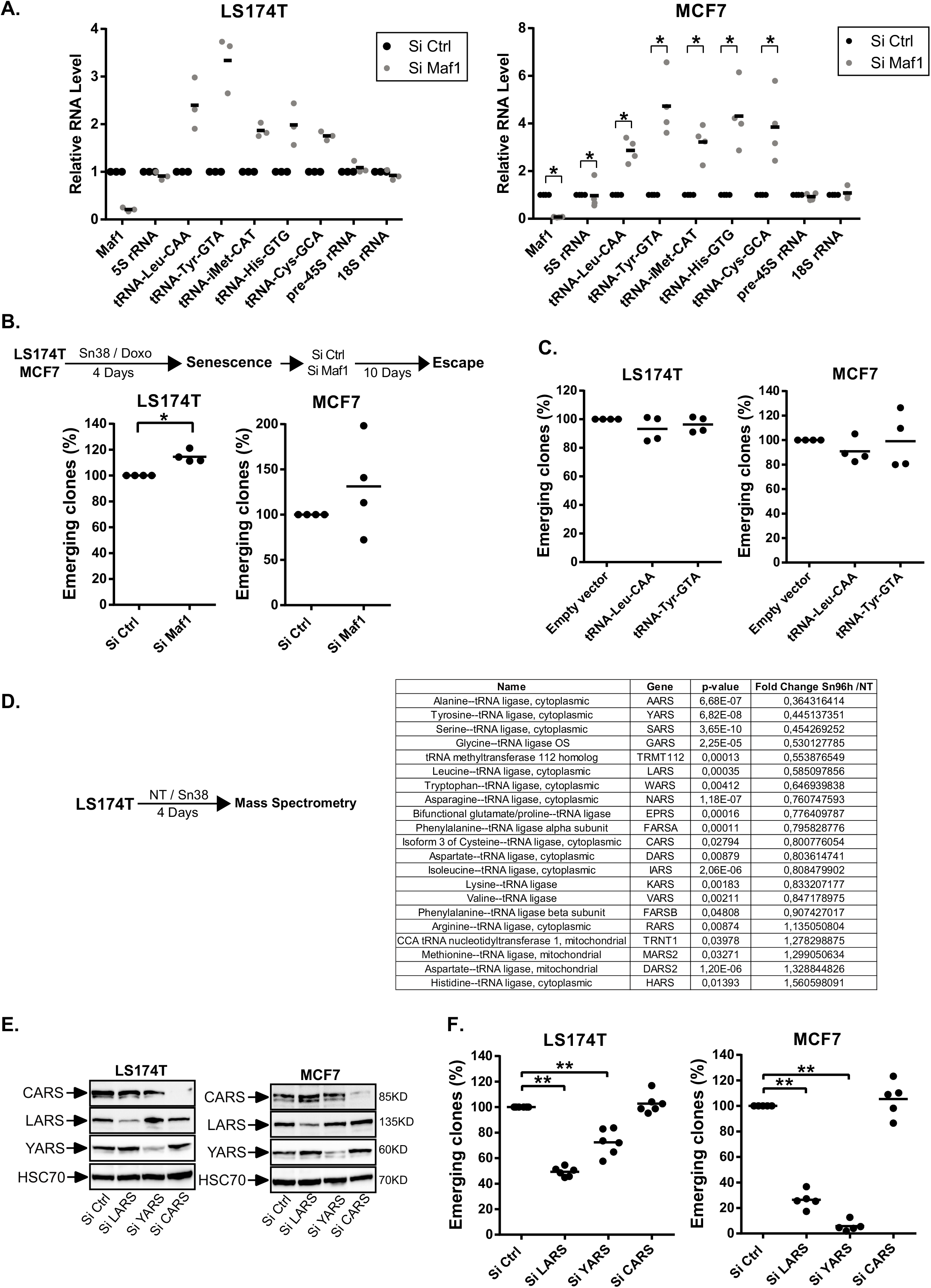
The LARS and YARS aminoacyl-tRNA synthetases are necessary for CIS escape. A. Senescent LS174T and MCF7 cells were transfected with a control siRNA or a siRNA directed against Maf1 during 24 hr and then washed with PBS and stimulated with fresh media. Cell extracts were recovered 2 days after Maf1 depletion and the expression of the indicated RNAs was analyzed by RT-QPCR (LS174T: n=3; MCF7: n=4, Kolmogorov-Smirnov test, * = p<0.05). B. Cells were treated and transfected as described above. Nine days after Maf1 inactivation, the number of emerging clones was analyzed (n=4, Kolmogorov-Smirnov test, * = p<0.05). C. LS174T and MCF7 senescent cells were transduced with an empty vector (pLKO.1) or a vector expressing the tRNA-Tyr-GTA or the tRNA-Leu-CAA. Ten days after, the number of emerging clones was analyzed (n=4). D. Quantitative proteomic analysis of proteins involved in tRNA biogenesis between growing and LS174T senescent cells (n=3). E. Senescent LS174T and MCF7 cells were transfected with a control siRNA or a siRNA directed against LARS, YARS or CARS. The depletion of the specifics Aminoacyl-tRNA Synthetases was validated by western blot 2 days after the transfection (n=2). F. Senescent LS174T and MCF7 cells were transfected with a control siRNA or a siRNA directed against LARS, YARS or CARS for 24 hr. The number of emergent clones was evaluated 9 days later (LS174T n=6, MCF7 n=5, Kolmogorov-Smirnov test, **= p<0.01).

We then determined if specific tRNAs could induce CIS escape. To this end, we infected senescent cells with lentivirus expressing the tRNA^Leu^-CAA and the tRNA^Tyr^-GTA. Results presented Figure 6C indicated that this did not modify the number of persistent clones (see also supplementary Figure 2E). This could be explained since endogenous tRNAs are probably expressed in large excess. Instead of increasing their expression, we made the hypothesis that tRNA activity might be regulated through the modulation of aminoacyl-tRNA synthetases. These enzymes allow the ligation of amino acid to their compatible tRNAs. We first used mass spectrometry to analyze their expression in our experimental conditions (Figure 6D). We observed that a significant number of tRNA ligases were down-regulated during senescence whereas others were up-regulated, indicating that this suppressive arrest did not induce a general inhibition of these proteins. To indirectly test the role of the tRNA^Leu^-CAA, tRNA^Tyr^-GTA and tRNA^Cys^-GCA, we down-regulated the expression of the leucine (LARS), tyrosine (YARS) and cystein (CARS) ligases (Figure 6E). Results presented Figure 6F indicated that LARS and YARS inactivation reduced senescence escape, both in LS174T and MCF7 cells. No effect was seen when the CARS ligase was inhibited.

Altogether, these results indicate that LARS and YARS are necessary for senescence escape and that this is specific since CARS had no effect on cell emergence.

### LARS and YARS modulate E2F-1 proliferative and apoptotic targets

To understand how the tRNA ligases could affect cell emergence, we inactivated YARS in senescent cells and performed a proteomic analysis after two days of emergence. In MCF7 and LS174T cells, a significant signature of the Rb and E2F1 pathway was detected (Figure 7A). This observation was interesting since recent results have connected the deregulation of ribosome biology to the Rb-E2F pathway (31). To validate this proteomic analysis, we first used MCF7 cells and observed using RT-QPCR experiments that the inactivation of YARS led to a down-regulation of E2F proliferative targets after two days of emergence (Figure 7B, left). Western blot analysis indicated that cyclin A expression was also down-regulated (Figure 7B, right). Interestingly, the down-regulation of this tRNA ligase increased the proportion of cells in the G2/M phase of the cell cycle and reduced S phase entry (Figure 7C).

**Figure 7:**
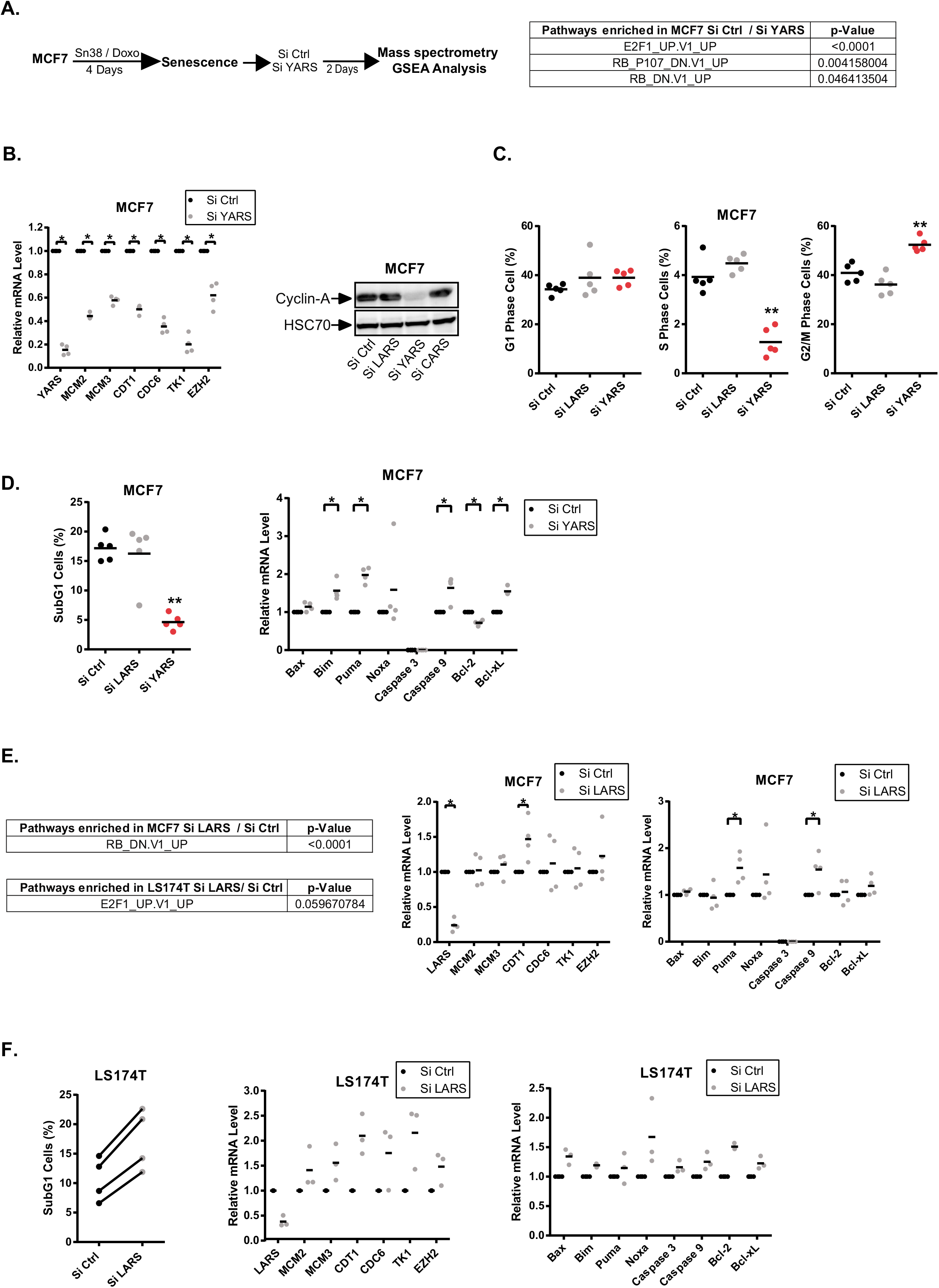
YARS and LARS regulate E2F1 targets during CIS escape. A. Senescent cells were transfected with a control siRNA or a siRNA directed against YARS. Cell extracts were analyzed by SWATH quantitative proteomics and GSEA analysis 2 days after depletion (n=3). B. Analysis by RT-QPCR and western blot of E2F1 proliferative targets in MCF7 cells following YARS depletion (n=4, Kolmogorov-Smirnov test, * = p<0.05, n=2 for western blot). C. Senescent cells were transfected with a control siRNA or a siRNA directed against YARS or LARS. After two days, FACS analysis was performed to analyze the cell cycle profile of the indicated cells (n=5, Kolmogorov-Smirnov test, ** = p<0.01). D. Senescent MCF7 cells were transfected as above, FACS analysis was performed to analyze apoptotic cells (n=5, Kolmogorov-Smirnov test, ** = p<0.01). The expression of the indicated mRNAs was analyzed in parallel by RT-QPCR (n=4, Kolmogorov-Smirnov test, * = p<0.05). E. MCF7 or LS174T senescent cells were transfected with a control siRNA or a siRNA directed against LARS. Cell extracts were analyzed by SWATH quantitative proteomics and GSEA analysis 2 days after depletion (n=3). In parallel, the indicated mRNAs were analyzed by RT-QPCR in MCF7 cells following LARS depletion (n=4, Kolmogorov-Smirnov test, * = p<0.05). F. LS174T senescent cells were transfected as above, FACS analysis was performed to analyze apoptotic cells (n=4). The expression of the indicated mRNAs was analyzed in parallel by RT-QPCR (n=3).

Besides cell cycle progression, E2F1 is a well-characterized activator of apoptosis. In MCF7 cells, YARS inhibition surprisingly reduced the percentage of apoptotic cells detected during the initial steps of senescence escape (Figure 7D). RT-QPCR experiments showed that pro-apoptotic members of the Bcl-2 family such as Puma or Bim were up-regulated as well as the Caspase 9 mRNA. All these genes have been previously described as E2F1 targets. However, these results also showed that Bcl-xL was up-regulated, suggesting that this survival protein inhibited its pro-apoptotic counterparts. Therefore, in MCF7, YARS inactivation prevented CIS escape and this was associated with a down-regulation of E2F1 proliferative targets. Note however that this was not observed in LS174T cells where these cell cycle and apoptotic targets were not modified (supplementary Figure 3A, 3B and 3C).

Following LARS inhibition, the GSEA analysis also detected a deregulation of the Rb-E2F1 pathway (Figure 7E). E2F1 proliferative targets were not down-regulated in MCF7 cells but an increased expression of Puma and Caspase 9 was detected when LARS was inhibited. In LS174T cells, an increased proportion of dying cells was observed and apoptotic genes such as Noxa were up-regulated (Figure 7F).

Altogether, these results indicate that LARS and YARS regulate the E2F1 pathway during senescence escape. However, this effect is heterogeneous and depends on the cell line, since apoptotic and proliferative genes were regulated differently by the two ligases and different observations were made in MCF7 and LS174T cells.

## DISCUSSION

Several studies have reported that senescence is much more dynamic than initially expected. How this heterogeneity is regulated remains largely unknown. It can be related to specific transcriptional activities, to the maintenance of epigenetic marks, to the variability of the SASP or to a specific oncogenic background in the case of cancer cells. Single-cell variability is expected to generate subpopulations of cells that enter distinct states of light or deep senescence and as a result some cells will escape chemotherapy more easily. In this study, we extend these observations and propose that the strength of CIS is related to the expression of specific tRNAs and aminoacyl-tRNA synthetases.

It is known that perturbations of ribosome biogenesis induce suppressive pathways. Ribosomal proteins such as L11 can interact with hdm2 to activate p53 and induce cell cycle arrest (32). Ribosomes are also controlled in a p53-independent manner since the RPS14 protein can interact with cdk4 to prevent its activity. This inhibits Rb phosphorylation and induces senescence as a response to an abnormal ribosome biogenesis (31). Our results indicate that tRNA transcription is inhibited during the initial step of the suppressive arrest but that emergent cells reactivate this activity to allow senescence escape. This effect is specific and not related to a general activation of RNA Pol III since only specific tRNAs are up-regulated during emergence. Of the aminoacyl-tRNA synthetases tested, only the LARS and YARS tRNA ligases allowed senescence escape. Importantly, the CARS ligase did not affect emergence and the expression of the corresponding tRNA^Cys^ was not modified. Given the complexity of tRNA biology, it remains to be determined on a larger scale if other tRNAs and ARSs are involved in cell emergence. We speculate that this will be the case and that different pools of tRNAs and ligases will regulate senescence pathways. Interestingly, recent results have shown that senescence prevents the ribosomal readthrough of stop codons and that cells that escape this suppressive mechanism have an abnormal translation termination (33). Accordingly, we speculate that a specific expression of tRNAs and ligases might regulate the expression of suppressive or oncogenic proteins. This could explain senescence heterogeneity and the ability of cells to remain definitely arrested or conversely to restart proliferation.

Further experiments are necessary to determine how tRNAs and ARSs regulate CIS and cell emergence. As described above (31), we have observed that these proteins deregulate the E2F1 pathway. Further experiments are now necessary to understand how YARS and LARS regulate E2F1 and if this is also mediated by RPS14 and cdk4 inhibition, but our results confirm that this signaling pathway is a main target of ribosome biology. It should also be noted that some aminoacyl-tRNA synthetases have adopted new functions during evolution, beyond protein synthesis. For instance, the leucyl-tRNA synthetase can induce mTOR activity upon leucine stimulation (34). The Trp-tRNA synthetase can interact with DNA-PK and PARP to activate p53 (35). It has also been recently reported that the seryl tRNA synthetase interacts with the POT1 member of the telomeric shelterin complex and that this accelerates replicative senescence (36). Thus, we can speculate that the LARS and YARS tRNA ligases have acquired new functions that somehow allow senescence escape.

New results have also described unexpected functions for tRNAs (37,38). Recent findings show that tRNA^Glu^ and tRNA^Arg^ are over-expressed in metastatic cells and that their inactivation reduces *in vivo* metastasis (39). These tRNAs induce a profound change in the proteome, with enrichment of proteins containing the GAA and GAG codons in the corresponding mRNA. This effect is specific, as it is not seen with other tRNAs. Santos et al. have recently reported that mutant tRNAs that mis-incorporate Serine instead of Alanine induce cell transformation (40). It has been proposed that a change in the pool of available tRNAs can lead to statistical errors in tRNA load within the ribosomal subunits (37,38). This generates a random proteome known as a statistical proteome, which represents mutated proteins or proteins with a novel sequence not completely coded for by the genome (41,42). It will be interesting to determine if the variability of the SASP might be explained by the utilization of different tRNA pools. Although this remains to be demonstrated, it leads to the hypothesis that subpopulations of senescent cells might express specific tRNAs (or tRNA ligases) and that this will allow the expression of secretomes presenting distinct, specific inflammatory activities.

Further experiments are therefore needed to determine if different pools of tRNAs and aminoacyl-tRNA synthetases are expressed in response to oncogenic insults, chemotherapy or during aging. We speculate that each type of suppressive arrest might lead to the expression of specific tRNAs and ARSs and that this might explain the specificity of these responses. In addition to epigenetic and transcriptional regulations, we therefore propose that the heterogeneity of tRNAs and ligases expression also leads to distinct states of light or deep senescence. This observation provides a rationale to further study the different facets of senescence responses and their links with chemotherapy resistance.

### Experimental procedures

See also the supplementary file

### Cell lines, senescence induction and generation of persistent cells

LS174T and MCF7 cell lines were obtained from the American Type Culture Collection. Cell lines were authenticated by STR profiling and were regularly tested to exclude mycoplasma contamination. To induce senescence, cell lines were treated for 96 hr in RPMI medium containing 3% of SVF with Sn38 (5 ng/ml, LS174T) or Doxorubicin (25 ng/ml, MCF7). To promote senescence escape cells were washed with PBS and stimulated with fresh medium containing 10% SVF for 7 (RT-qPCR analysis, Western Blot, SA-β galactosidase staining) or 10 days (evaluation of emerging clone number).

### Treatments

Cells were treated with the following drugs: Torin-1 (Cell Signaling, 14379): 15 nM, Rapamycin (Santa-Cruz, sc-3504): 5 nM, Tunicamycin (Santa-Cruz, sc-3506): 0.1 to 8 µg/ml, Thapsigargin (Santa-Cruz, sc-24017): 10 to 50 nM

### SiRNA transfection

Cells were transfected with 50 nM of small interfering RNA against p21(CDKN1A) (ON-TARGET plus Human CDKN1A (1026) Dharmacon, L-00341-00-0005), Maf1 (ON-TARGET plus Human MAF1, Dharmacon, L-018603-01-0005), BRF1 (ON-TARGET plus human BRF1, Dharmacon, L-017422-00-0005), LARS (ON-TARGETplus Human LARS siRNA, L-010171-00-0005), YARS (ON-TARGETplus Human YARS siRNA, L-011498-00-0005), CARS (ON-TARGETplus Human CARS siRNA, L-010335-01-0005) and prevalidated control siRNA (Dharmacon, D-001810-10-20) using DharmaFect-4 (Dharmacon, T-2004-03). Note that the siRNA concentration was reduced to 12,5nM for the tRNA ligases to reduce cell toxicity.

### ShRNA, lentiviruses and cell transduction

pLKO.1-TSC2 was a gift from Do-Hyung Kim (Addgene plasmid #15478), pLKO-TRC2 (scramble ShRNA, Sigma mission, SH216). For the generation of lentiviruses, 293 cells were cotransfected with with the packaging plasmids (pMDLg/pRRE, pRSV-Rev, and PMD2.G) and the different PLKO plasmids by lipofectamine 2000 for 24 hr. After 24 hr, the medium was replaced with fresh medium. After 48 hr, virus-containing supernatant was collected and centrifuged for 5 minutes at 300g and filtered through 0.45 μm. For the transduction, 2.5 ml were used to transduce LS174T and MCF7 senescent cells in the presence of 4 µg/ml Polybrene (Santa Cruz). After 24h, the media was replaced. For tRNA over-expression, the TRY-GTA5-3 tRNA-Tyr and TRL-CAA2-1 tRNA-Leu sequence were cloned in the pLKO.1 vector.

### Cell Cycle Analysis

250 000 cells were incubated with 150 µL of solution A (trypsin 30 µg/mL, Sigma) for 10 min at room temperature in the dark. Then, 125µL of solution B (trypsin inhibitor 0,5 mg/mL, RNAse A 0,1 mg/mL, Sigma) was added for 10 min in the dark. Finally, cells were incubated with 125 µL of solution C (propidium iodide 0,6 mM, spermine tetrahydrochloride 3,3 mM, Sigma) for 10 min at 4°C. All the solutions were prepared in a storage buffer pH 7,6 containing 3,4 mM sodium citrate 2H_2_0 (Sigma), 0,1% Igepal CA-630 (Sigma), 3 mM spermine tetrahydrochloride and 1mM de tris-aminomethane.

### Patient-derived organoids

After informed consent, breast tumors from patients who underwent surgical tumor resection at the ICO Paul Papin Cancer Center were processed through a combination of mechanical disruption and enzymatic digestion by collagenase to generate patient-derived organoids (PDO) as described (30). Briefly, isolated cells were plated in adherent basement membrane extract drops (BME type 2, R&D systems, 3533-010-02) and overlaid with optimized breast cancer organoid culture medium. Medium was changed every 4 days and organoids were passaged every 1-3 weeks. Organoids were treated by 2 cycles of doxorubicin (25 ng/ml) for 96 hrs. At the end of the first cycle, the medium was changed and organoids were incubated 3 days without chemotherapy before the second cycle. Following the 2 cycles, the medium was replaced and changed every 3-4 days to analyze chemotherapy escape.

## Acknowledgements

This work was supported by grants from the Ligue Contre le Cancer (Comité du Maine et Loire, du Finistère, de la Loire Atlantique), the Rotary Club (Maine et Loire) and the SIRIC ILIAD Nantes-Angers-INCA-DGOS-Inserm grant 12558

## Abbreviations

CIS: Chemotherapy-Induced Senescence
ARS: aminoacyl-tRNA synthetases

## Supplementary Figures

**Supplementary Figure 1.**
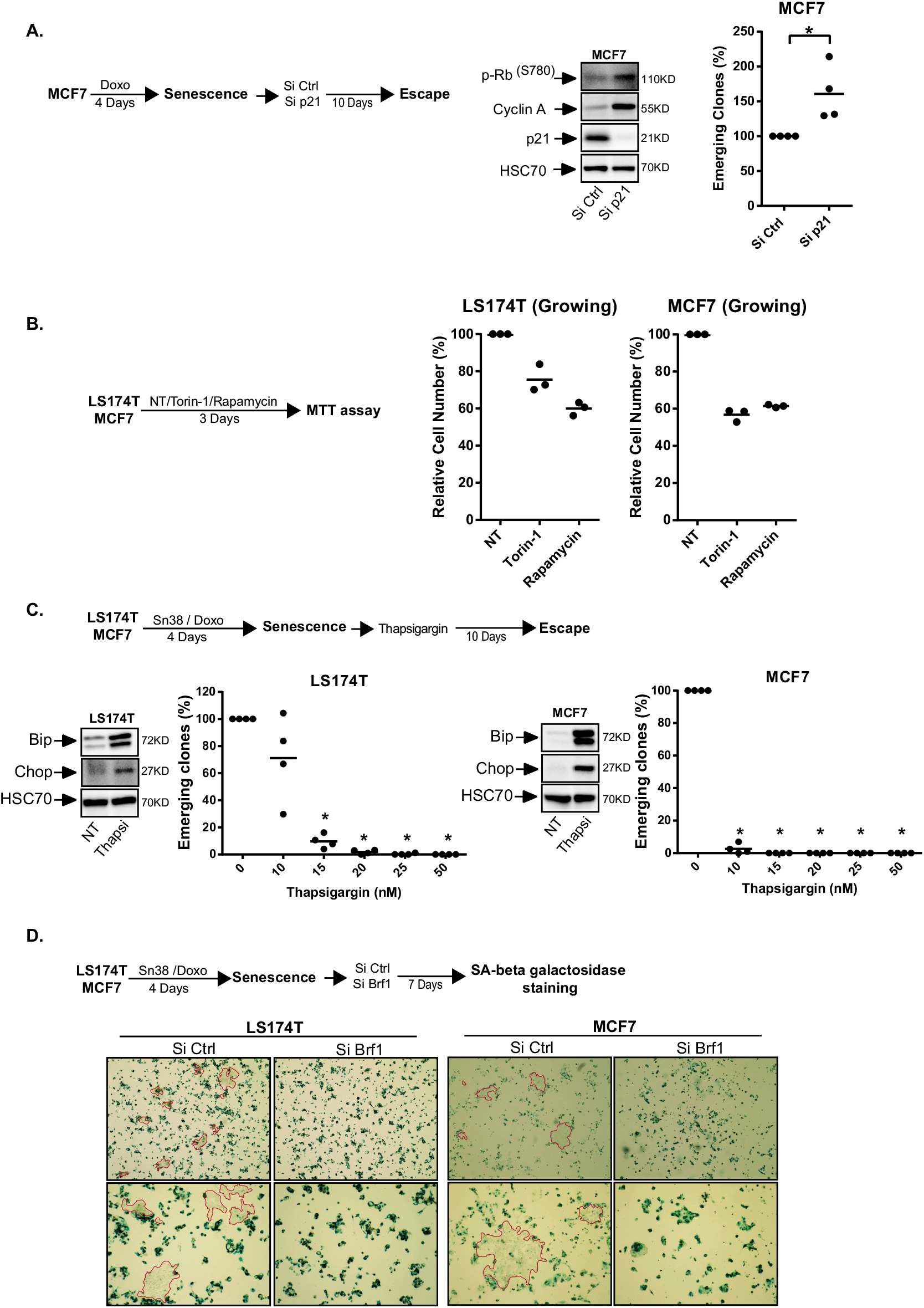
**A**. Senescent MCF7 cells were transfected with a control siRNA or a siRNA directed against p21 for 24 h and persistent cells were then generated by adding 10% FBS. p21 down-regulation as well as the expression of cyclin A and Rb phosphorylation were evaluated by Western blot two days after the depletion. Emergence was evaluated after 10 days (n=4, Kolmogorov-Smirnov test, * = p<0.05, n=3 for western blots). **B**. LS174T and MCF7 cells were treated or not with mTOR inhibitors (Torin-1: 15nM; Rapamycin: 5nM), and cell viability was analyzed by MTT assay after 3 days. (n=3). **C**. Senescent cells were treated with increasing concentrations of thapsigargin and the number of emerging clones was evaluated 10 days later (n=4, Kolmogorov-Smirnov test,* = p<0.05). The induction of UPR was validated by Western blot after 24h using a 10 nM concentration (n=2). **D**. Representative images of SA-β galactosidase staining 7 days after BRF1 inactivation in LS174T and MCF7 senescent cells. Growing persistent cells are underlined in red (n=3).

**Supplementary Figure 2.**
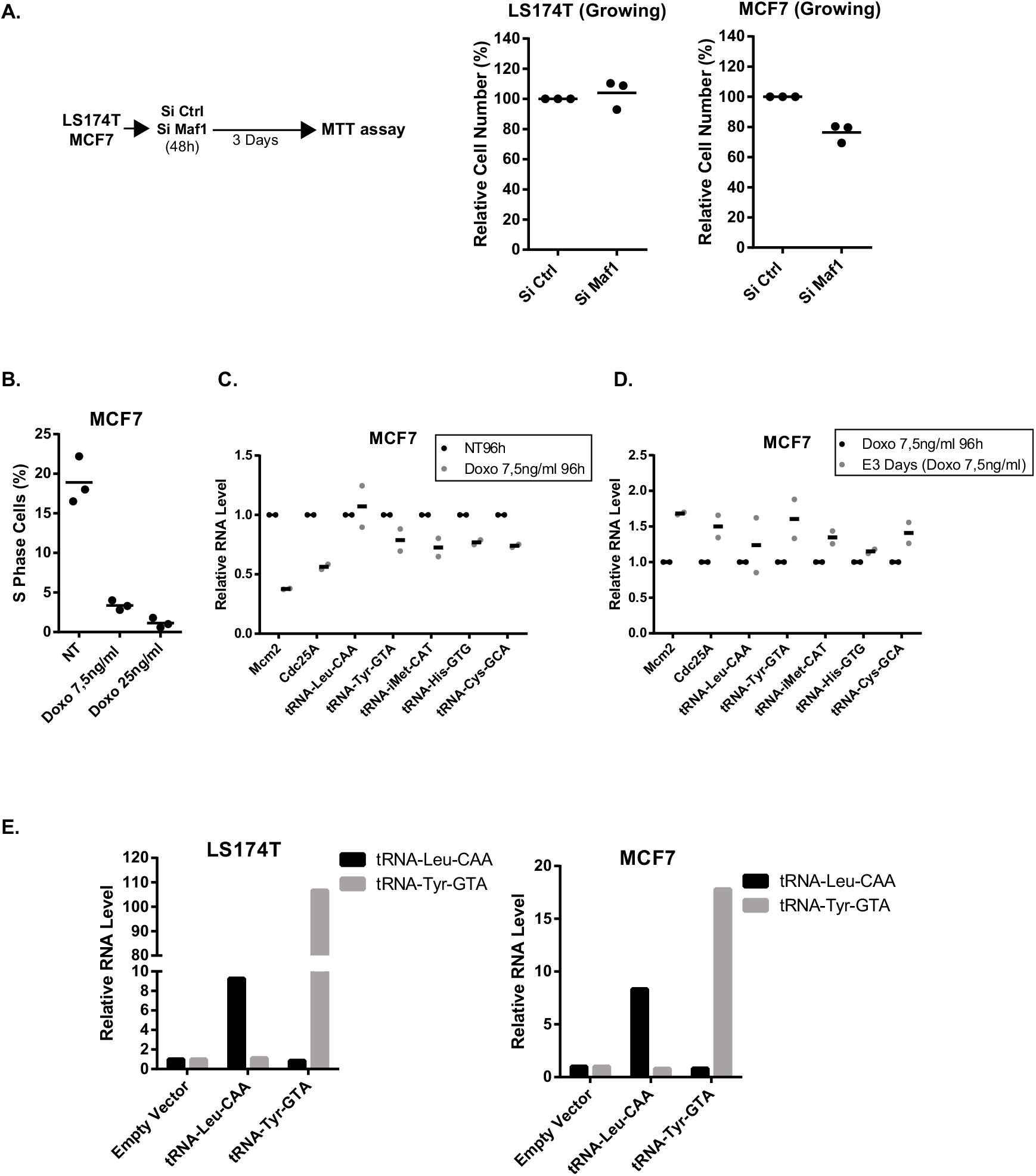
**A** LS174T and MCF7 cells were transfected with a control siRNA or a si RNA directed against Maf1 for 48h. Cell viability was analyzed by MTT assay after 3 days (n=3). **B**. MCF7 cells were treated or not with 7,5ng/ml or 25ng/ml of Doxorubicin during 4 days. FACS analysis was then performed to analyze the percentage of cells in S phase (n=3). **C**. RT-QPCR anlaysis of the indicted proliferative markers (Mcm2 and Cdc25a) and tRNAs in MCF7 cells treated or not with Doxorubicin (7,5ng/ml) during 4 days (n=2). (**D**) RT-QPCR analysis of the expression of the indicated RNAs in MCF7 cells. Cells have been treated for 4 days with Doxorubicin (7,5ng/ml) and emergence was induced during 3 days by fresh media addition (E3, n=2). **E**. LS174T and MCF7 senescent cells were transduced with an empty vector (pLKO.1) or a vector expressing tRNA-Tyr-GTA or tRNA-Leu-CAA. 10% FBS was then added for two days and RNA expression was analyzed by RT-QPCR.

**Supplementary Figure 3.**
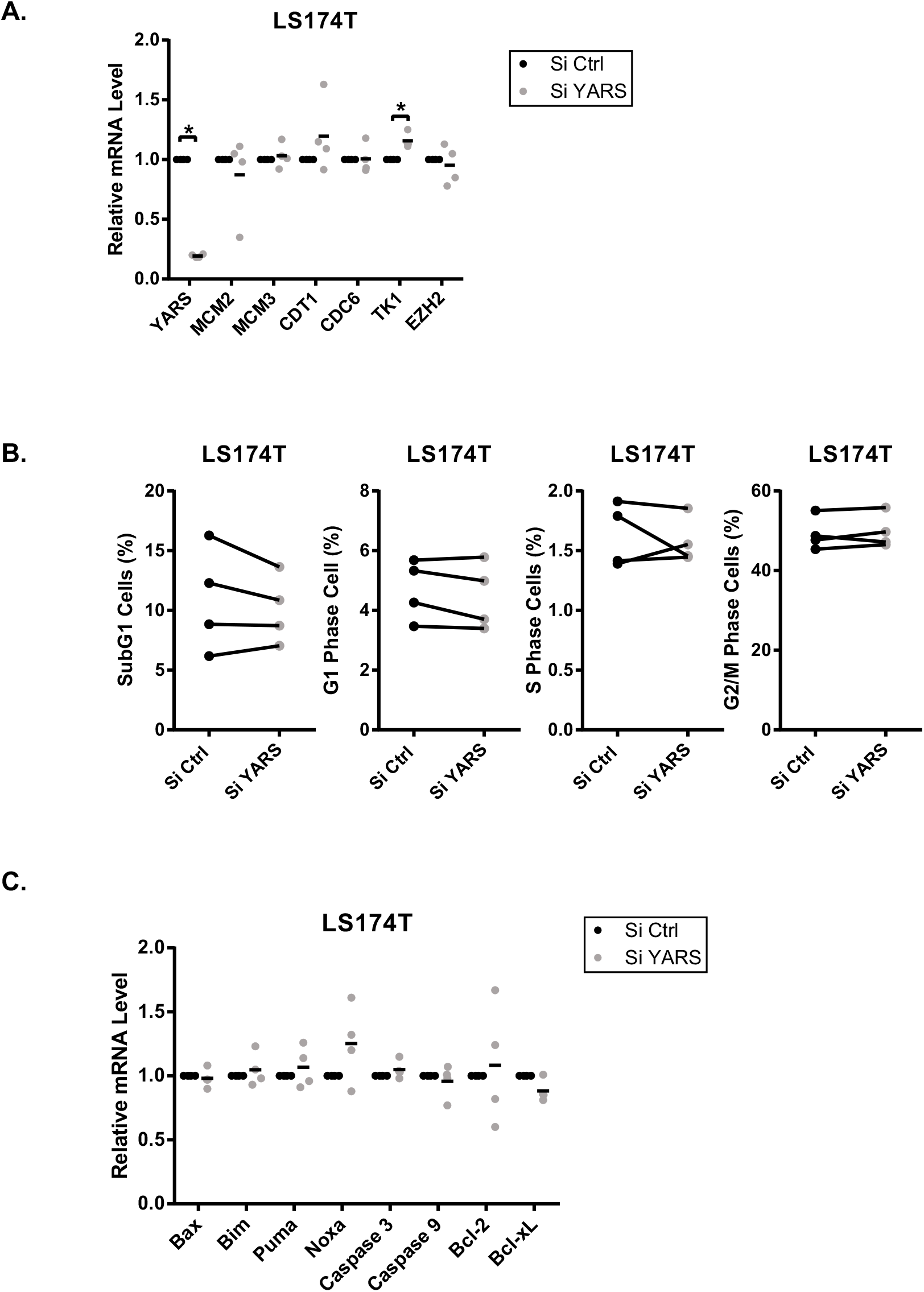
**A**. LS174T senescent cells were transfected with a control siRNA or a siRNA directed against YARS. Two days after the depletion, the expression of E2F1 proliferative targets was analyzed by RT-QPCR (n=4, Kolmogorov-Smirnov test,* = p<0.05). **B**. LS174T senescent cells were transfected with a control siRNA or directed against YARS. Two days after the depletion, FACS analysis was performed to analyze the cell cycle profile of the indicated cells (n=4). **C**. LS74T senescent cells were transfected with a control siRNA control or directed against YARS. Two days after the depletion, RT-QPCR was performed to analyse the expression of the indicated mRNAs (n=4).

## Supplementary Methods

### Chromatin immunoprecipitation (Chip)

Cells were cross-linked with 1% formaldehyde (Sigma Aldrich) for 10 min at room temperature. Cross-linking was stopped by adding 0.125 mol/L glycine for 5 min. Cells were washed three times with cold phosphate-buffered saline (PBS). Cells were then scraped and washed three times with cold PBS. Pellets were resuspended in 1 mL of lysis buffer (5 mM PIPES, 85mM KCl, and 0.5% NP40). All buffers were supplemented with proteases and phosphatase inhibitors (1mM PMSF, 10 µg/ml aprotinin, 10 µg/ml leupeptin, 10 µg/ml pepstatin, 1 mM Na3VO4, and 50 mM NaF). Samples were incubated for 15 min at 4 °C and vortexed for 30 seconds every 2 minutes. Cells were centrifuged for 10 min, 16 000g at 4 °C. Supernatants were discarded and pellets were resuspended in 500 µL of sonicating buffer (10 mM EDTA, 1% SDS, and 50 mM Tris-EDTA, pH 8). Nuclear extracts were sonicated (20 cycles of 23 s of sonication and 25s on ice) to obtain DNA reverse cross-linked fragments with a size of 500-200 bases. Supernatants were diluted 10 times with IP buffer (0.01% SDS, 1.1%Triton X-100, 1.2 mM EDTA, 16.7 mM Tris-HCl (pH =8.0), and 167 mM NaCl). For each condition, 13 µg of chromatin were used (chromatin concentration was estimated by nanodrop on the reverse cross-linked chromatin). The chromatin was pre-cleared with 25 µl of beads for 2 hr on rotation at 4°C. Beads (Protein A/G Magnetic Beads, Thermofisher, 26162) were coated during the same time with 3 µg of the following antibodies : Rabbit Polyclonal POLR3A (Abcam, ab96328) or Rabbit IgG, polyclonal - Isotype Control (Abcam, ab171870). Each pre-cleared sample was then incubated with 45 µl of coated magnetic beads and DTT (1 mM) and BSA (10 µg/ml) added at the last moment. After overnight incubation at 4°C, the beads were washed successively with 1.5 ml of TSE1 Buffer (1%Triton X-100, 150 mM NaCl, 20 mM Tris-HCl, pH 8.1, 0.1% SDS, and 2 mM EDTA), 1.5mL of TSE2 buffer (1%Triton X-100, 500 mM NaCl, 20 mM Tris-HCl, pH 8.1,0.1% SDS, and 2 mM EDTA), and 1.5mL of TSE3 buffer 1% NP40, 1% sodium deoxycholate, 250mM LiCl, and 10mM Tris-HCl, pH 8.1). Following two washes in TE buffer (10 mM Tris-HCl and 1 mM EDTA), samples were eluted with 300 µL of fresh elution buffer (1% SDS and 0.1M NaHCO3). The cross-link was reversed by adding 24µL of NaCl (5 M) and 6µL of EDTA (0.5 M) to the samples and incubating overnight at 65 °C. DNA was purified using a High Pure PCR Template Preparation Kit (Roche) and analyzed by Q-PCR using 4 µl of the elution and 6 µl of Syber green containing the primers (5 µl of Syber green and 1 µl of primers (5 µM)).

### Mass spectrometry

The technical approach was the same as previously reported (Guillon, Petit et al. 2019).

#### Creation of the spectral library

In order to build the spectral library, peptide solutions of several protein samples were analyzed by a shotgun approach by micro-LC–MS/MS. Five pooled samples of breast, colorectal and blood tissues were prepared to obtain a good representation of the peptides. Each sample was fractionated by offgel fractionator in 24 fractions. Each fraction was separated into a micro-LC system Ekspert nLC400 (Eksigent, Dublin, CA, USA) using a ChromXP C18CL column (0.3 mm × 15 cm, 3 µm, 120 Å) (Eksigent) at a flow rate of 5 µL/min. Water and ACN, both containing 0.1% formic acid, were used as solvents A and B, respectively. The following gradient of solvent B was used: 0 to 5 min 5% B, 5 to 125 min 5% to 35% B, then 9 min at 95% B, and finally 9 min at 5% B for column equilibration. As the peptides eluted, they were directly injected into a hybrid quadrupole-TOF mass spectrometer Triple TOF 5600 + (Sciex, Redwood City, CA, USA) operated with a ‘top 30’ data-dependent acquisition system using positive ion mode. The acquisition mode consisted of a 250 ms survey MS scan from 400 to 1250 m/z, followed by an MS/MS scan from 200 to 1500 m/z (75 ms acquisition time, 350 mDa mass tolerance, rolling collision energy) of the top 30 precursor ions from the survey scan. The peptide and protein identifications were performed using Protein Pilot software (version 5.0, Sciex) with a human Swiss-Prot/TrEMBL concatenated target-reverse decoy database (downloaded in March 2016) containing 142,441 target human protein sequences, specifying MMTS as Cys alkylation. The false discovery rate (FDR) was set to 0.01 for both peptides and proteins. The MS/MS spectra of the identified peptides were then used to generate the spectral library for SWATH peak extraction using the add-in for PeakView Software (version 2.2, Sciex) MS/MSALL with SWATH Acquisition MicroApp (version 2.0, Sciex). Peptides with a confidence score above 99% as obtained from Protein Pilot database search were included in the spectral library.

#### Relative quantification by SWATH acquisition

LS174T cells were analyzed using a DIA method. Each sample (5 µg) was analyzed using the LC– MS equipment and LC gradient described above, using a SWATH-MS acquisition method. The method consisted of repeating the whole gradient cycle, which consisted of the acquisition of 35 TOF MS/MS scans of overlapping sequential precursor isolation windows (25 m/z isolation width, 1 m/z overlap, high sensitivity mode) covering the 400 to 1250 m/z mass range, with a previous MS scan for each cycle. The accumulation time was 50 ms for the MS scan (from 400 to 1250 m/z) and 100 ms for the product ion scan (230 to 1500 m/z), thus making a 3.5 s total cycle time.

#### Data analysis

The targeted data extraction of the SWATH runs was performed by PeakView using the MS/ MSALL with SWATH Acquisition MicroApp. PeakView processed the data using the spectral library created from the shotgun data. Up to ten peptides per protein and seven fragments per peptide were selected, based on signal intensity; any shared and modified peptides were excluded from the extraction. The retention times from the peptides that were selected for each protein were realigned in each run according to iRT peptides (Biognosys AG, Schlieren/Zürich, Switzerland) spiked in each sample and eluting along the whole time axis; the extracted ion chromatograms were generated for each selected fragment ion. PeakView computed a score and an FDR for each assigned peptide using chromatographic and spectra components; only peptides with an FDR of less than 5% were used for protein quantitation. The peak areas for peptides were obtained by summing the peak areas of the corresponding fragment ions; protein quantitation was calculated by summing the peak areas of the corresponding peptides. MarkerView (version 1.2, Sciex) was used for signal normalization, and differential abundance was tested by applying a T-test at the protein level.

### Western Blot

Following cell lysis with FASP Buffer (0.1 M Tris-HCL, 4% SDS, pH=7.6) containing a cocktail of inhibitors (10 µg/ml aprotinin, 10 µg/ml leupeptin, 10 µg/ml pepstatin, 1 mM Na3VO4, 50 mM NaF), lysates were sonicated and then boiled for 10 min. Proteins were separated on a SDS polyacrylamide gel and transferred to a PVDF membrane. Following a 1 hr incubation in 5% milk, Tris-buffered saline (TBS), and 0.1% Tween 20, the membranes were incubated overnight at 4 °C with the following primary antibodies : p21waf1/cip1 (1/1000, Cell Signaling 2947), p-S6 ribosomal (S235/236) (1/1000, Cell Signaling, 2211), Chop (1/1000, Cell Signaling, 5554), HSC70 (1/1000, Santa Cruz, sc-7298), Bip (1/1000, Cell Signaling, 3177), p-Rb(S780) (1/1000, BD Pharmingen, 558385), Cyclin A (1/1000, Santa Cruz, sc-271682), TSC2 (1/1000, Cell Signaling, 4308), LARS (1/1000, Cell Signaling, 35509), TyrRS (1/1000, Santa-Cruz, sc-166741), CysRS (1/1000, Santa-Cruz, sc-390230). Membranes were then washed three times with TBS with 0.1% Tween 20 and incubated for 45 minutes with the secondary antibodies listed below: Anti-rabbit IgG, HRP-linked antibody (1/3000, Cell Signaling, 7074), Anti-mouse IgG, HRP-linked Antibody (1/3000, Cell Signaling, 7076). Visualization was performed by chemiluminescence with a Fusion Solo (Vilber).

### RT-QPCR Analysis

Analysis was performed using the comparative CT method (2^(ΔCt)), according to the expression of three endogenous housekeeping gene TBP, PPIA and EEF1A1. All primers sequences are provided below.

### SA-β Galactosidase staining

Cells were fixed for 10 min at room temperature in 2% formaldehyde, washed with PBS and incubated at 37°C in the absence of CO_2_ with freshly-made staining solution: 0.3 mg/mL of 5-bromo-4-chloro-3-indolyl-β-d-galactopyranoside)(X-Gal,Promega, V394A), 40 mM citric acid (Sigma), 40 mM sodium phosphate (Sigma) (stock solution (400 mM citric acid, 400 mM sodium phosphate) must be at pH 6), 5 mM potassium hexacyanoferrate (Sigma), 5 mM potassium ferricyanide (Sigma), 150 mM NaCl (Sigma), 150 mM MgCl2 (Sigma). SA-β galactosidase staining was observed after 16 hours.

### MTT assay

Cell viability and proliferation was evaluated by the MTT (3-4,5-dimethylthiazol 2,5-diphenyltetrazolium bromide) assay. Cells were seeded in 96-well clear-bottomed plates at a density of 2000 cells (LS174T) and 900 cells (MCF7) in RPMI medium containing 10% SVF. After 72 hr, 40 µL of 5 mg/ mL MTT solution was added to each well and incubated in a humidified 5% CO_2_ atmosphere at 37°C for 3 hours. After incubation, the cells were pelleted and dried. Next, 100 µL of DMSO (dimethylsulfoxyde) were added to each well, and the cells were incubated at room temperature for 1 hr. Absorbance was measured by a microtiter plate reader at 562 nm.

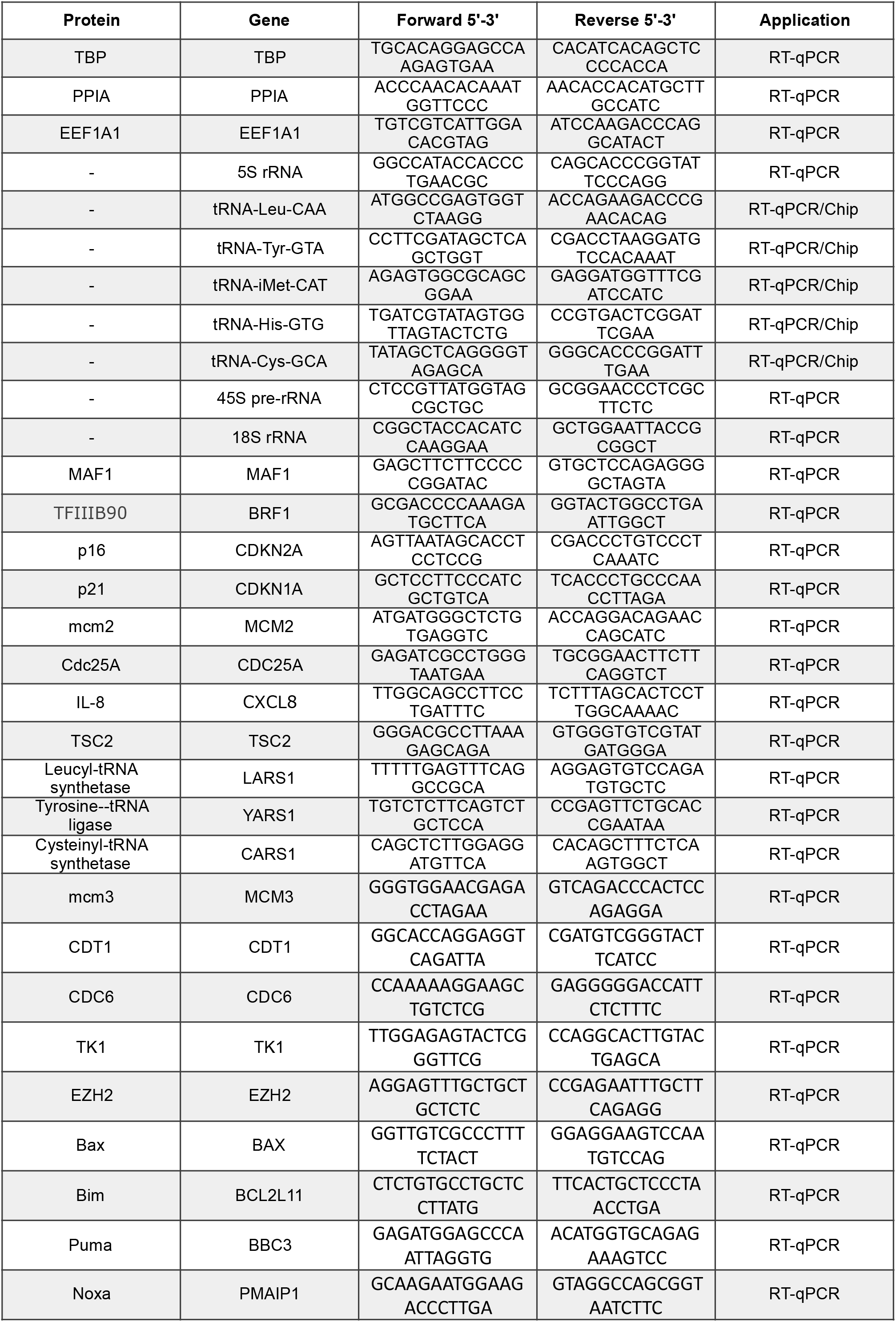
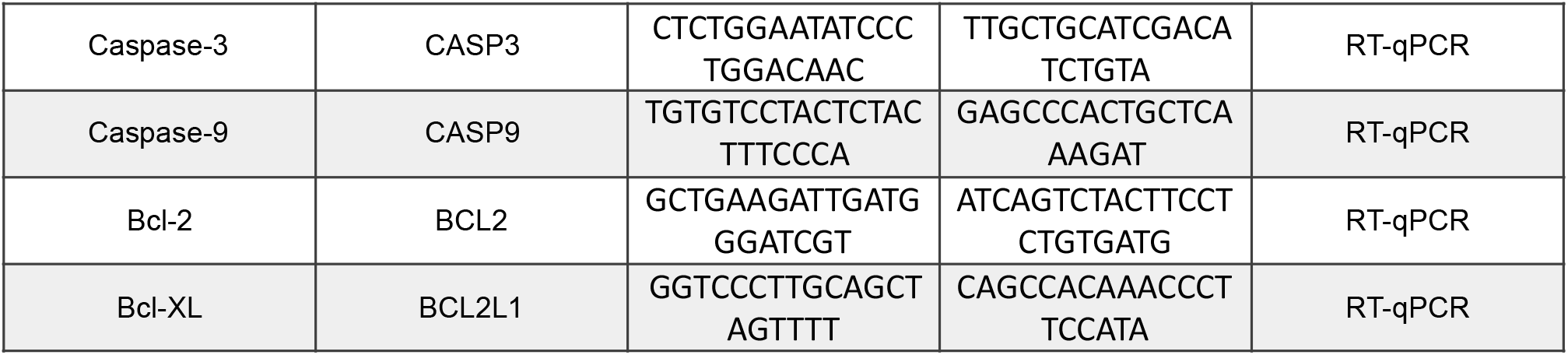

